## Supplementary Information for "Investigating the origin and consequences of endogenous default options in repeated economic choices"

***Supplementary Experiments***

This section describes two preliminary experiments for the experiments in the main text. Analogous to the main text, preliminary Experiment S1 refers to a gain task and preliminary Experiment S2 to a loss task.

### Methods

#### Participants

Both preliminary experiments were approved by the local Ethics Committee of the University of Amsterdam. All participants gave informed consent prior to taking part in the study. The participants were recruited from the laboratory's participant database.

31 participants enrolled in Experiment S1 (gain version: 25 females, mean age = 24.8 SD = 8.9; 29) and 30 in Experiment S2 (loss version); targeted sample size = 30. However, one participant was excluded from the analyses in Experiment S2 for always choosing the option displayed on one side of the screen, leaving 29 participants in the reported analyses (22 females, mean age = 22.1, SD = 4.4). Participants were paid according to a performance-based payment incentive, rather than a purely flat-fee payment [1]. In Experiment S1, participants had the opportunity to choose between a course credit (1 ECTS) or a financial compensation (base amount of 8€), with the possibility of getting an extra amount of 4€, depending on a randomly chosen trial. In Experiment S2, they had the opportunity to get a financial compensation (base amount of 12€), with a potential subtraction from this compensation of 4€, depending on a randomly chosen trial. The conversion rates between experimental euros (EE)/real euros (€) were set individually, such that the highest outcome lottery faced by each participant scaled to a 4 real € gain (Experiment S1) or loss (Experiment S2). This led to a conversion rate of approximatively 20 EE/€ in Experiment S1, and 25 EE/€ in Experiment S2.

#### Design

Both preliminary experiments were programmed in Cogent for Matlab®. We designed two repeated binary decision-making tasks involving probabilistic monetary outcomes, one framed as potential gains (Experiment S1) and the other as potential losses (Experiment S2) (**Supplementary Figure 1A** and **B**). At each trial, participants were presented with a wheel-of-fortune which defined two options. In Experiment S1, the option featured as the Safe lottery offered a probability $p$ > 50% (e.g., 75%) of winning a certain amount of money $a$ (e.g., 4€), and the option featured as the Risky lottery offered a probability $1-p$ (e.g., 25%) to win a higher amount $A$ (e.g., 8€). In Experiment S2, the option featured as the Safe lottery offered a probability $p$ > 50% (e.g., 75%) of losing a certain amount of money $a$ (e.g., 4€), and the option featured as the Risky lottery offered a probability $1-p$ (e.g., 25%) to lose a higher amount $A$ (e.g., 8€). The probabilities were presented as the two complementary areas of the wheel-of-fortune and the amounts as vertical bars of varying height. In Experiment S1, the bars were up and in Experiment S2, they were down relative to the center of the screen. The attributes of the risky and safe lotteries were colored in blue or yellow in both preliminary experiments, and this color attribution was balanced across participants. The side of presentation (left or right) of the safe and risky lotteries was also balanced across trials. Text describing exact probabilities and amounts was presented on the same screen. In Experiment S2, amounts were displayed with a minus sign.

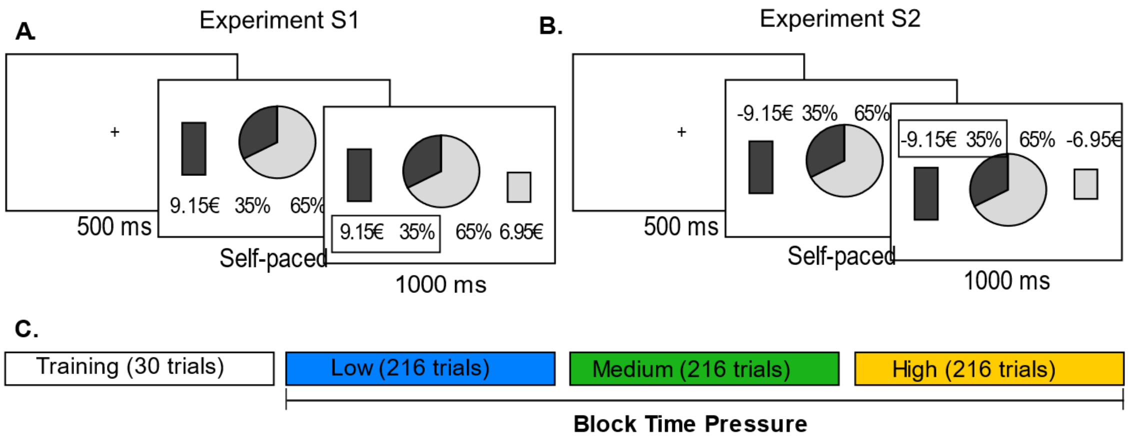

**Supplementary Figure 1**. **Preliminary Experiments.** Behavioral tasks of Experiment S1 and S2 (**A.** and **B.**, respectively). Successive screenshots displayed during a given trial are illustrated from left to right, with durations in milliseconds. At each trial, following a short fixation (500ms), subjects have to choose between a risky (here, left: 35% chance to win or lose 9.15€) and a safe (here, right: 65% chance to win or lose 6.95€) lottery. Choices are self-paced, and followed by a choice-confirmation screen, where the selected lottery is highlighted by a contour box. **C.** Experimental design. Subjects start with a training session to familiarize themselves with the task. In the experimental session (3 even blocks of 216 trials), three conditions are implemented, with increasing time pressure.

Both preliminary experiments involved two phases – a training session and an experimental session (**Supplementary Figure 1C**). During the training session, participants experienced 30 trials, to familiarize with the task and the probabilities: once participants selected their preferred option, the wheel-of-fortune was spun, and participants could win – in the case of the gain task – or lose – in the case of the loss task – the amount paired with the chosen option if the spin stopped on the chosen-option portion. After the pointer was rotated, explicit feedback was given (e.g., "You win *a* €!" or "You lose *a* €!", respectively). In the experimental session, participants performed 3 blocks of 216 trials, in a self-paced mode. The experimental task operated under the same principle of the training task, except that the spinning and feedback were not visible on the computer screen. The absence of feedback was mainly to avoid explicit learning effects. We induced different levels of time pressure across the 3 blocks: in Block 1, participants were instructed to be "as thoughtful as possible" in their choices. In Block 2, to be "as thoughtful and as fast as possible". Finally, in Block 3, to be "as fast as possible". At the end of the experiment, one of the participants' choices was selected, executed, and the resulting gains/losses were added/subtracted to their payoffs. The instructions and payoffs were thus designed to induce increasing time pressure across the blocks, while still incentivizing preference-based choices. All the payment procedures were explained in detail before the experiment.

### Stimuli

The stimulus set was generated so that 1) all lottery pairs would be unique and 2) the set of lottery pairs would span a fixed distribution of choice difficulty.
In each lottery pair, there was always one safe lottery with high probability of winning or losing a low amount (Experiment S1 and S2, respectively) and a risky lottery with low probability of winning or losing a high amount.

Let $a$ and $P$ be the amount and probability of the safe lottery, and $A$ and $p$ the amount and probability of the risky lottery. We set the following:

(Eq. 1)

$$0.5 < P < 1$$

(Eq. 2)

$$p = 1 - P$$

(Eq. 3)

$$a > 0$$

(Eq. 4)

$$A > 0$$

(Eq. 5)

$$a < A$$

We proceeded as follows: we choose vectors of $P$ and $a$, which satisfied certain conditions, given the desired vector of choice difficulty that we aimed for (see below). We then determined $A$, so that choice trials would exhibit desired properties. Specifically, we aimed at generating lottery pairs with fixed distributions of expected value ($EV$) difference between the safe and risky options ($\Delta EV$).

(Eq. 6)

$$\Delta EV = EV_{safe} - EV_{risky}$$

In Experiment S1, we proceeded as follows.
We choose 6 values of $P$ and 6 values of $\Delta EV$, which remained constant over blocks:

(Eq. 7)

$$P \in [0.55\ 0.60\ 0.65\ 0.70\ 0.80\ 0.85];$$

(Eq. 8)

$$\Delta EV \in [-1 - 0.75 - 0.5 - 0.25\ 0.25\ 0.5];$$

Note that the distribution of $\Delta EV$ is not symmetric, and biased towards negative values. In other words, the risky lotteries have on average bigger expected values than the safe lotteries. This was designed to counterbalance the natural tendency of subjects to be risk averse in the gain domain, and make sure that they would end up choosing a significant proportion of risky lotteries.
Then, for each time pressure block (216 trials), we defined a new set of 6 values for $a$. This defined a quasi-factorial design with 6 ($a$) × 6 ($P$) × 6 ($\Delta EV$) = 216 trials per block. Note that once $\Delta EV$, $P$ and $a$ have been defined, the value of $A$ can be uniquely derived as follows:

Let us start with our chosen definition of choice difficulty, namely $\Delta EV$

(Eq. 9)

$$\Delta EV = P \times a - (1 - P) \times A$$

Let us set
(Eq. 10)

$$A = a + d$$

with
(Eq. 11)

$$d > 0.$$

In this way, Eq. 3, 10 and 11ensure that conditions specified in Eq. 4) and 5) are satisfied. We could write, then, that:
(Eq. 12)

$$\Delta EV = P \times a - (1 - P) \times (a + d)$$

so that:
(Eq. 13)

$$d = \frac{a(2P - 1) - \Delta EV}{1 - P} > 0,$$

and
(Eq. 14)

$$a > \frac{\Delta EV}{2P - 1}$$

If $\Delta EV < 0$, (Eq. 14) is already implied by condition (Eq. 3), so there is no extra constraint on $a$. However, if $\Delta EV > 0$, (Eq. 14) there is an extra constraint on $a$, such that that
(Eq. 15)

$$a > \max\left(\frac{\Delta EV}{2P - 1}\right)$$

In our case, it means that we had to define vectors of a, with values greater than 5; Note that we defined a new set of 6 values for $a$ at each block, hence generating a unique set of lottery pairs per block.
At each trial, knowing $a$, $P$ and $\Delta EV$, we can derive $A$:
(Eq. 16)

$$A = a + \frac{a\ (2p - 1) - \Delta EV}{1 - P}$$

Note that all amounts were rounded to the 5 cent (experimental euro) precision.
The stimulus set was generated in the same way for Experiment S1 and S2, but the distribution of choice difficulty as specified in (Eq.8) was reversed for Experiment S2. Specifically, while the vector of $\Delta EV$ was biased towards the risky option in Experiment S1(see above), it was biased towards the safe option in Experiment S2. This reversed distribution was applied so as to counterbalance risk-seeking behavior in the loss domain.

### Statistical Analysis

Similar generalized linear mixed models (*GLMMs*) were applied to both choices and RTs in both preliminary experiments. GLMMs were performed with the lmerTest package [2], an extension of lme4 package [3] in R. We used a linear link function when modelling RTs, and a binomial (logit) link function when modelling choices. Choices were coded as 1 for safe and 0 for risky. Time pressure (TP, three levels: Low, Medium, High) was coded as cardinal – i.e., as continuous, linear variables (taking values -1, 0 and 1 for increasing levels of TP).
In Wilkinson-Rogers notation, GLMMs take the form *y ~ fixed + (random)*, where *y* corresponds to the dependent variable, *fixed* to the independent variables used to explain or predict the dependent variable, and *random* to the deviations from the predicted values due to subjects' variability. Only one model was tested in both preliminary experiments, with time pressure as an independent variable. The formulas of the fixed effects were, then, *choice ~ time pressure* and *rt ~time pressure*, for choices and RTs, respectively. Random-effects structure included individual random intercepts and slopes, namely (1 + *time pressure* | *subjects*). These account for differing baseline-levels of choices and reaction times (the intercept, represented by 1) as well as differing responses to the main factor(s) in question, that is "time pressure".

### Model Parameter Estimates

In the next tables, we display the estimated fixed-effect coefficients of the GLMMs accounting for choices and RTs in Experiment S1 and S2.

| Coefficients | Experiment S1 | Experiment S2 |
|---|---|---|
| **Intercept** | $\beta$ = 1.22539; SE = .23148;<br>z = 5.294; p < .001 | $\beta$ = -1.7121; SE = .2797;<br>z = -6.122; p < .001 |
| **TP** | $\beta$ = .24144; SE = .09082;<br>z = 2.658; p = .00785 | $\beta$ = -.2231; SE = .0622;<br>z = -3.588; p < .001 |

**Supplementary Table 1.** Estimated coefficients of the models accounting for choices in Experiment S1 and S2. TP: time pressure.

| Coefficients | Experiment S1 | Experiment S2 |
|---|---|---|
| **Intercept** | $\beta$ = 2157.6; SE = 233.1;<br>df = 31.0; t = 9.257; p <.001 | $\beta$ = 1840.0; SE = 156.6;<br>df = 28.0; t = 11.748; p <.001 |
| **TP** | $\beta$ = -1320.5; SE = 174.8;<br>df = 31.0; t = -7.553; p <.001 | $\beta$ = -1225.7; SE = 158.4;<br>df = 28.0; t = -7.737; p <.001 |

**Supplementary Table 2.** Estimated coefficients of the models accounting for RTs in Experiment S1 and S2. TP: time pressure.

**Results**

A first manipulation control analysis revealed that participants successfully decreased their RTs under increasing time pressure in both preliminary experiments (Experiment S1: $\beta_{TP}$ = -1320.50, SE = 174.80, p < .001; Experiment S2: $\beta_{TP}$ = -1225.70, SE = 158.40, p < .001). In addition, we found that participants also modulated their choice pattern under increasing time pressure so as to increase the proportion of safe choices in Experiment S1, and of risky choices in Experiment S2 (Experiment S1: $\beta_{TP}$= .24, SE = .09, p = .00785; Experiment S2: $\beta_{TP}$ = -.22, SE = .06, p < .001 – see Supplementary **Figure 2A** and **B**).

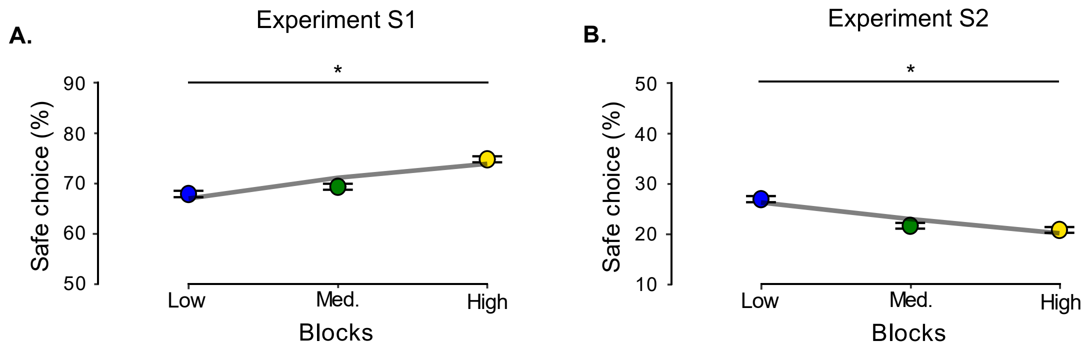

**Supplementary Figure 2. Results Preliminary Experiments.** Observed proportion of safe choices in Experiment S1 and S2 (**A.** and **B.**, respectively). Results show that subjects increasingly chose the safer option in Experiment S1, and the risky option in Experiment S2, under increasing time pressure. Dots and error bars indicate sample mean and within-subject standard error of the mean. Lines represent the average fit of the generalized linear mixed-effects regressions.

### ***Supplementary Methods***

This section provides supplementary information on Experiment 1 and 2 in the main text.

#### Additional Behavioral Predictions

This section extends on the behavioral predictions introduced in the main text.

An important contribution of this study is the definition of several potential dimensions of default options. We have derived different predictions on the pattern of risky vs. safe choices observed in response to the manipulation of time pressure, time on task, and choice proportion, which we attributed to the presence of natural, dominant, and learned default options (see **Main text – Figure 1**).

Here, we further consider additional factors which can also create systematic patterns of risky vs. safe choices in response to the manipulation of time pressure, time on task, and choice proportion.

Because those patterns are not consistent with the results of the preliminary Experiment S1 and S2 (see section **Supplementary Experiments**), however, we did not include those in our main default option hypotheses.

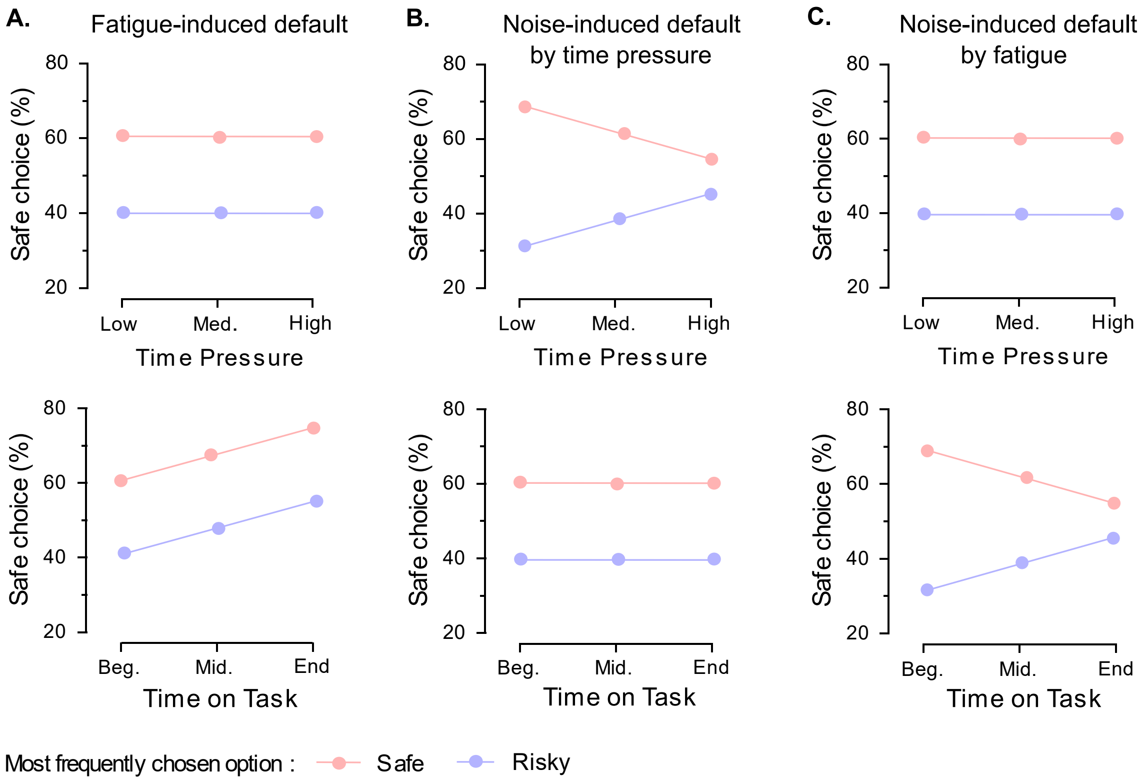

**Supplementary Figure 3. Hypotheses about other potential default options. A.** Fatigue-induced default. In this scenario, with increasing time on task, individuals gradually loose the cognitive resources to make decisions, and therefore increasingly choose - and stick to - a preferred option, regardless of the manipulation of choice proportion [4,5]. **B.** Noise induced by time pressure. In this scenario, individuals' choices become increasingly noisy/random as time pressure increases. This pattern is consistent with changes in response boundaries in a sequential sampling framework, as often argued in the context of perceptual choices [6,7]. **C.** Noise induced by fatigue. In this last scenario, individuals' choices become increasingly noisy as time on task increases. This could be typically due to the accumulation of fatigue or boredom, decreasing attention or cognitive effort that individuals are willing to invest to render accurate decisions [8,9].

### Stimuli Generation

This section describes the procedure used to generate the lotteries in Experiment 1 and 2 in the main text (gain and loss tasks).
Similar to the preliminary experiments, the stimulus set was generated so that 1) all lottery pairs would be unique and 2) the set of lottery pairs would span a fixed distribution of choice difficulty.
In each lottery pair, there was always one safe lottery with high probability/low amount and a risky lottery with low probability/high amount.
Let $a$ and $P$ be the amount and probability of the safe lottery, and $A$ and $p$ the amount and probability of the risky lottery. We set (Eq. 1), (Eq. 2), (Eq. 3), (Eq. 4) and (Eq. 5) – see section **Supplementary Experiments**.
And we proceeded as follows: we choose vectors of $P$ and $a$, given the desired vector of choice difficulty that we aimed for (see below). We then determined $A$, so that choice trials would exhibit expected utility between lotteries, not expected value as for the preliminary experiments.

### Model fitting

In the preliminary experiments, stimuli were generated using the difference in expected value between the two lotteries as a proxy for difficulty. Yet, our results (see **Supplementary Experiments**) show that participants are significantly risk-averse when facing gain lotteries and risk-seeking when facing loss lotteries. To better control for these risk attitudes, and generate the stimuli for Experiment 1 and 2, we therefore fitted an expected utility model [10,11] to the data of preliminary experiments (S1 and S2, respectively), so as to estimate the parameters best capturing our participants' choices.

A common practice is to model individual's choices as a soft-maximization of expected utility, using a logistic choice function. Defining $p_s$ as the probability of choosing the safe lottery, we have:

(Eq. 17)

$$p_s = 1/(1 + \exp(-\boldsymbol{\beta} \times \Delta EU)$$

where

(Eq. 18)

$$\Delta EU = EU_{safe} - EU_{risky} = P \times a^{\mathbf{u}} - (1-P) \times A^{\mathbf{u}}.$$

This model has two free-parameters, $\boldsymbol{\beta}$ and $\mathbf{u}$, which respectively represent the inverse temperature (i.e. how stochastic vs. utility maximizing choices are), and the utility curvature (i.e., how risk-seeking vs. risk-averse choices are). Note that the utility curvature $\mathbf{u}$ is often considered a proxy for risk preference, given that $\mathbf{u} > 0$ will typically generate risk-seeking choices, while $\mathbf{u} < 0$ typically generate risk-averse choices [11].

Model parameters were estimated at the individual level by minimizing free energy [12,13], using a variational Bayes approach under the Laplace approximation [14]. In order to avoid any bias in the parameter estimation, we only used data from the no time pressure block (216 choices). We used pretty wide, unbiased priors (mean (±SD): $\boldsymbol{\beta} = 3\ (10)$; $\mathbf{r} = 1\ (10)$).

Results (mean( ± STD)) of the estimation procedure were for Experiment 1: **β** = 4.3804 (3.6485) and **r** = 0.5592 (0.5400); and for Experiment 2: **β** = 2.3660 (2.2043) and **r** = 0.7492 (0.3651).

### Modelling Parameter Recovery

To check the quality of the parameter estimation procedure, we performed a parameter recovery analysis: we simulated 100 synthetic participants, with $\boldsymbol{\beta}$ and $\mathbf{r}$ respectively sampled from a gamma distribution (shape = 1.2, scale = 5) and from a normal distribution (mean = 1; standard deviation = 0.5).

We then estimated the correlation between parameters estimated from the VBA scheme outline above, and the parameters used to generate the synthetic data (**Supplementary Figure 4**).

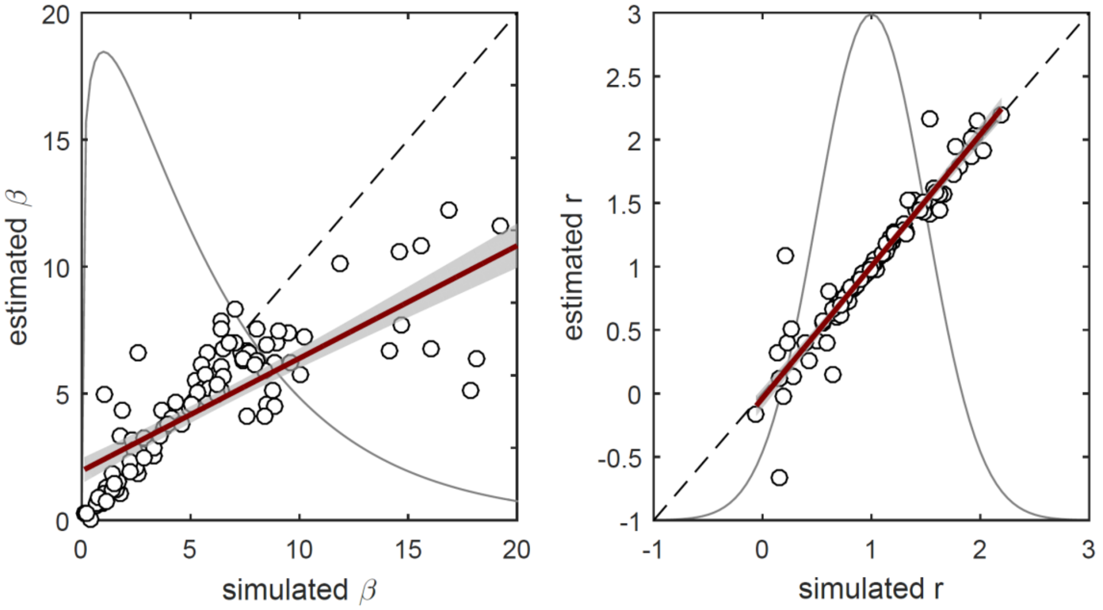

**Supplementary Figure 4. Parameter recovery.** Overall, data from 100 synthetic participants were simulated. The 2 estimated parameters (left: **β**; right: **r**) per participants were then regressed against the true parameters used for simulating the data. Results show very good identifiability – although high values of **β** seem to be slightly underestimated. Each dot represents a synthetic individual. The red continuous lines the best linear fits, and the shaded grey areas the 95% confidence interval around the best-linear fit. The grey densities represent the probability distributions used to sample the parameters.

### Model-based Stimuli Generation

In Experiment 1 and 2 the set of lottery pairs was designed to offer a tighter experimental control over individual's probability of choosing one class of lottery (safe or risky), under the assumption that their decisions could be reasonably described by the model and averaged parameters (**β** and/or **r** ) estimated in the preliminary experiments S1 and S2, respectively.
In what follows, we describe the procedure to generate the stimuli set in Experiment 1, and for the safe condition. At the end of this section, more details can be found about how this procedure was adapted to generate stimuli in Experiment 2, and for the risky condition.
We choose 6 values of $P$ and 9 values of $p_s$:

(Eq. 19)

$$P \in [0.55\ 0.60\ 0.65\ 0.70\ 0.75\ 0.80];$$

(Eq. 20)

$$p_s \in [.50\ .55\ .60\ .65\ .70\ .75\ .80\ .85\ .90];$$

We also fixed 3 values of $a$ and 3 values of $A$:
(Eq. 21)

$$a \in [3\ 5\ 10];$$

(Eq. 22)

$$A \in [40\ 50\ 60];$$

This defined a quasi-factorial design with 9 ($p_s$) × 6 ($P$) × 6 ($3a + 3A$) = 324 trials. Note that once $p_s$, $P$ and $a$ (or $A$) have been defined, the value of $A$ (or $a$) can be uniquely derived as follows. Rearranging (Eq. 17), we get

(Eq. 23)

$$\Delta EU = \frac{1}{\boldsymbol{\beta}} \times log(\frac{p_s}{1 - p_s})$$

Combined with (Eq. 18), this gives

(Eq. 24)

$$P \times a^{\mathbf{u}} - (1 - P) \times A^{\mathbf{u}} = \frac{1}{\boldsymbol{\beta}} \times log(\frac{p_s}{1 - p_s}),$$

so that,

(Eq. 25)

$$a = (\frac{\frac{1}{\boldsymbol{\beta}} \times \log(\frac{p_s}{1 - p_s}) - (1 - P) \times A^{\mathbf{u}}}{P})^{1/\mathbf{u}}$$

and

(Eq. 26)

$$A = (\frac{\frac{1}{\boldsymbol{\beta}} \times \log(\frac{p_s}{1 - p_s}) + P \times a^{\mathbf{u}}}{1 - P})^{1/\mathbf{u}}$$

The 324 trials were divided in 18 blocks of 18 trials, such that each 18-trial block consisted of 2 repetitions of each $p_s$. This ensured that the distribution of choice probability was constant over blocks, and averaged at 70%. Note that initial values of $a$ and $A$ (Eq. 21-22) were handpicked to satisfy all necessary conditions (Eq. 1-5). Despite our careful control of the experiment setting, we noticed that our choice of $a$ in Experiment 1 - safe condition was misguided, such that 18 trials did not satisfy Eq 5 (i.e. displayed lotteries with A < a). Yet, excluding those trials from our analyses did not change any of the reported results.
For Experiment 1/ risky condition, the setup was similar. Except that we took

$$p_s \in [.50\ .45\ .40\ .35\ .30\ .25\ .20\ .15\ .10];$$

For Experiment 2, we used parameters estimated from the preliminary Experiment S2 (loss task). The procedure was similar, but the value of $A$ and $a$ was adapted as follows:

$$A = [-40 \ -50 \ -60];$$

$$a = [-20 \ -25 \ -30];$$

A similar procedure was used to generate the 72 stimuli for the framing session. We used the following parameters:

$$P \in [.55\ .60\ .65\ .70\ .75\ .80];$$

In addition, for Experiment 1, we defined:

$$p_s \in [.60\ .75\ .90];$$

$$A = [45\ 65];$$

$$a = [4\ 7.5];$$

For Experiment 2, we defined:

$$p_s \in [.10\ .25\ .40];$$

$$A = [-45 - 65];$$

$$a = [-22 - 27];$$

This resulted in a quasi-factorial design with 3 ($p_s$) × 6 ($P$) × 4 ($2a + 2A$) = 72 trials.

### Generalized Linear Mixed Models (GLMMs)

This section describes the GLMMs used to test the effects of time pressure, time on task, and choice proportion on choices and response times (RTs) in Experiment 1 and 2.

#### Models Specification

Similar *GLMM*s were applied to both choices and RTs. In the following tables, we specify the formulas of the fixed-effects (left side of the tables), which varied iteratively until a winning model was found (bold model in the right side) – i.e. when removing an interaction and/or variable did significantly decrease the goodness-of-fit of the model.
Random-effects structure was kept fixed across all models, and included individual random intercepts and slopes, namely (1 + *time pressure* + *time on task* | *subjects*). These account for differing baseline-levels of choices and reaction times (the intercept, represented by 1) as well as differing responses to the main factor(s) in question, that is "time pressure and time on task".

We ran additional models, with no individual random intercepts, so that individual baseline differences would not interfere with the estimation of our between-subject factor. However, because we found no difference with the results reported in the tables (estimated with random intercepts), we do not report them.

| Sequence of models | Experiment 1 | Experiment 2 |
|---|---|---|
| CH ~ TP * ToT * CP | AIC = 10377; BIC = 10480; *LL* = -5174.4; $\chi^2$= .5236; df = 1; p = .4693 | AIC = 9775.9; BIC = 9879.0; *LL* = -4874.0; $\chi^2$= .2216; df = 1; p = .6378 |
| CH ~ TP + ToT + CP + ToT : CP + TP : CP + TP : ToT | AIC = 10375; BIC = 10471; *LL* = -5174.6; $\chi^2$= .0427; df = 1; p = .8363 | AIC = 9774.2; BIC = 9869.9; *LL* = -4874.1; $\chi^2$= .0756; df = 1; p = .7833 |
| CH ~ TP + ToT + CP + ToT : CP + TP : ToT | AIC = 10373; BIC = 10462; *LL* = -5174.7; $\chi^2$= .6804; df = 1; p = .4094 | AIC = 9772.2; BIC = 9860.6; *LL* = -4874.1; $\chi^2$= 1.5244; df = 1; p = .217 |
| **CH ~ TP + ToT + CP + ToT : CP** | **AIC = 10372; BIC = 10453; *LL* = -5175.0; $\chi^2$= 6.9157; df = 1; p = .008544** | **AIC = 9771.8; BIC = 9852.8; *LL* =** -4874.9; **$\chi^2$= 6.3222; df = 1; p = .01192** |
| CH ~ TP + ToT + CP | AIC = 10377; BIC = 10451; *LL* = -5178.5 | AIC = 9776.1; BIC = 9849.7; *LL* = -4878.0 |

**Supplementary Table 3.** Sequential model comparison of GLMMs accounting for choices. All models used a logit link-function (logistic regression). Left column: structure of the fixed-effects, indicated in Wilkinson-Rogers notation. This table includes model diagnostics for Experiment 1(middle column) and Experiment 2 (right column) task. From top to bottom, models were iteratively derived from the full model (first line), by deleting the least significant parameter. At each step, a likelihood-ratio test was performed, to assess if the simplified model was significantly decreasing the model goodness-of-fit. The procedure was stopped when this test came significant. AIC: Akaike information criterion. BIC: AIC information criterion. LL: log likelihood. $\chi^2$: chi-square statistics of the likelihood-ratio test between current model and next simpler model (one line below). df: degree of freedom of the likelihood-ratio test. p: p-value of the likelihood-ratio test. CH: choices. TP: time pressure. ToT: time on task. CP: choice proportion.
Bold font indicates the winning model from the iterative procedure. Note that in both cases, the winning model also has the lowest AIC.

| Sequence of models | Experiment 1 | Experiment 2 |
|---|---|---|
| **RT ~ TP * ToT * CP** | **AIC = 208312; BIC = 208423; *LL* = -104141; $\chi^2$= 10.6180; df = 1; p = .00112** | AIC = 220625; BIC = 220735; *LL* = -110297; $\chi^2$= .9360; df = 1; p = .3333 |
| RT ~ TP + ToT + CP + TP : ToT + TP : CP + ToT : CP | AIC = 208321; BIC = 208424; *LL* = -104146 | AIC = 220624; BIC = 220727; *LL* = -110298; $\chi^2$= $.2^{-4}$ ; df = 1; p = .9899 |
| RT ~ TP + ToT + CP + TP : ToT + ToT : CP | - | AIC = 220622; BIC = 220718; *LL* = -110298; $\chi^2$= .2284; df = 1; p = .6327 |
| **RT ~ TP + ToT + CP + TP : ToT** | - | **AIC = 220620; BIC = 220709; *LL* = -110298; $\chi^2$= 180.2109; df = 1; p <.001** |
| RT ~ TP + ToT + CP | - | AIC = 220798; BIC = 220879; *LL* = -110388 |

**Supplementary Table 4.** Sequential model comparison of GLMMs accounting for RTs . All models used an identity link-function (linear regression). Left column: structure of the fixed-effects, indicated in Wilkinson-Rogers notation. This table include model diagnostics for Experiment 1(middle column) and Experiment 2 (right column) task. From top to bottom, models were iteratively derived from the full model (first line), by deleting the least significant parameter. At each step, a likelihood-ratio test was performed, to assess if the simplified model was significantly decreasing the model goodness-of-fit. The procedure was stopped when this test came significant. AIC: Akaike information criterion. BIC: AIC information criterion. LL: log likelihood. $\chi^2$: chi-square statistics of the likelihood-ratio test between current model and next simpler model (one line below). df: degree of freedom of the likelihood-ratio test. p: p-value of the likelihood-ratio test. RT: reaction-times. TP: time pressure. ToT: time on task. CP: choice proportion.
Bold font indicates the winning model from the iterative procedure. Note that in both cases, the winning model also has the lowest AIC.

### Model Parameter Estimates

In the next tables, we display the estimated fixed-effect coefficients of the GLMMs accounting for choices and RTs in Experiment 1 and 2.

| Coefficients | Experiment 1 | Experiment 2 |
|---|---|---|
| **Intercept** | β = 1.09287; SE = .44916; z = 2.433; p = .014969 | β = -1.75167; SE = .50983; z = -3.436; p < .001 |
| **TP** | β = .22103; SE = .08669; z = 2.550; p = .010781 | β = -.28152; SE = .08851; z = -3.181; p = .001469 |
| **ToT** | β = .14819; SE = .07827; z = 1.893; p = .058333 | β = -.36337; SE = .13916; z = -2.611; p = .009023 |
| **CP** | β = -2.20177; SE = .63284; z = -3.479; p < .001 | β = 1.72600; SE = .57999; z = 2.976; p = .002921 |
| **ToT : CP** | β = -.30842; SE = .11837; z = -2.606; p = .009173 | β = .46292; SE = .18115; z = 2.555; p = .010607 |

**Supplementary Table 5.** Estimated coefficients of the winning models accounting for choices in Experiment 1 and 2. TP: time pressure. ToT: time on task. CP: choice proportion.

| Coefficients | Experiment 1 | Experiment 2 |
|---|---|---|
| **Intercept** | β = 1937.11; SE = 242.97; df = 37.02; t = 7.973; p <.001 | β = 2775.82; SE = 272.58; df = 46.04; t = 10.183; p <.001 |
| **TP** | β = -1074.02; SE = 201.42; df = 37.04; t = -5.332; p <.001 | β = -2077.32; SE = 232.10; df = 36.00; t = -8.950; p <.001 |
| **ToT** | β = -151.24; SE = 105.40; df = 37.20; t = -1.435; p = .159642 | β = -614.81; SE = 159.07; df = 36.00; t = -3.865; p <.001 |
| **CP** | β = 431.33; SE = 348.32; df = 37.00; t = 1.238; p = .223396 | β = 246.28; SE = 218.57; df = 36.00; t = 1.127; p = .267310 |
| **TP : ToT** | β = 124.75; SE = 33.59; df = 11612.07; t = 3.714; p <.001 | β = 570.85; SE = 42.36; df = 11556.00; t = 13.477; p <.001 |
| **TP : CP** | β = -334.42; SE = 288.72; df= 37.01; t = -1.158; p = .254164 | - |
| **ToT : CP** | β = -187.79; SE = 150.99; df = 37.08; t = -1.244; p = .221422 | - |
| **TP : ToT : CP** | β = 155.19; SE = 47.61; df = 11611.61; t = 3.259; p = .001120 | - |

**Supplementary Table 6.** Estimated coefficients of the winning models accounting for RTs in Experiment 1 and 2. TP: time pressure. ToT: time on task. CP: choice proportion.

### *Supplementary References*

1. Brase G. How different types of participant payoffs alter task performance. Judgm Decis Mak. 2009;4.
